# Angiosperm speciation speeds up near the poles

**DOI:** 10.1101/619064

**Authors:** J. Igea, A. J. Tanentzap

**Affiliations:** Department of Plant Sciences, University of Cambridge, Downing St, Cambridge, CB2 3EA, UK

**Keywords:** latitudinal diversity gradient, speciation, angiosperms, macroevolution, biogeography, biodiversity

## Abstract

Recent evidence has questioned whether the Latitudinal Diversity Gradient (LDG), whereby species richness increases towards the Equator, results from higher rates of speciation in the tropics. Allowing for time heterogeneity in speciation rate estimates for over 60,000 angiosperm species, we found that the LDG does not arise from variation in speciation rates because lineages do not speciate faster in the tropics. These results were consistently retrieved using two other methods to test the association between occupancy of tropical habitats and speciation rates. Our speciation rate estimates were robust to the effects of both undescribed species and missing taxa. Overall, our results show that speciation rates follow an opposite pattern to global variation in species richness. Greater ecological opportunity in the temperate zones, stemming from less saturated communities, higher species turnover or greater environmental change, may ultimately explain these results.

## Introduction

Biodiversity on Earth is very unevenly distributed. The Latitudinal Diversity Gradient (LDG), whereby species richness increases towards the Equator, is the most prominent example of this unevenness. This pervasive pattern has fascinated biologists for more than two centuries (von Humboldt 1808; Wallace 1878; Pianka 1966; Hawkins 2001), but its underlying causes remain largely unknown.

A major class of explanations proposes that elevated rates of speciation generate the higher diversity in the tropics and so primarily create the LDG (Mittelbach *et al*. 2007). This increase in speciation is generally attributed to higher environmental energy in the tropics, which in turn can hasten evolution through shorter generation times and higher mutation rates (Dowle *et al*. 2013), and is known as the “evolutionary speed hypothesis” (Rensch 1959; Rohde 1992). Other explanations propose that elevated speciation rates near the Equator can stem from higher chances of allopatric speciation caused by narrower thermal niches (Ghalambor *et al*. 2006), or from stronger biotic interactions (e.g., faster coevolution) – the ‘Red Queen running faster when she is hot’ (Brown 2014). However, while some studies on mammals (Rolland *et al*. 2014), amphibians (Pyron & Wiens 2013) and butterflies (Condamine *et al*. 2012) have shown that tropical lineages speciate faster, other studies have found no latitudinal differences (Rabosky & Huang 2015; Economo *et al*. 2018) or higher rates of speciation in temperate areas (Weir & Schluter 2007), particularly when focusing on recent rates of speciation (Schluter 2016; Schluter & Pennell 2017; Rabosky *et al*. 2018). Recent speciation rates should be less affected by extinction (Nee *et al*. 1994) and more accurate than deep-time estimates (Louca & Pennell 2019), even in the absence of paleontological evidence (Marshall 2017). They should also reflect any variable that historically influenced speciation and also varied latitudinally, such as environmental energy (Title & Rabosky 2019).

Flowering plants are one of the largest eukaryotic radiations with roughly 350,000 species and show a marked LDG (Francis & Currie 2003). Despite this pattern, latitudinal differences in macroevolutionary rates in angiosperms have received comparatively little attention at a species-level. Previous efforts have shown that angiosperm diversification is faster in tropical families (Jansson & Davies 2008) and in tropical lineages within a limited number of clades (Svenning *et al*. 2008; Jansson *et al*. 2013). The only species-level analysis found no differences in speciation, but did so by using a method that assumed that all rate differences in the angiosperm Tree of Life (ToL) were due to geographic distribution alone (Antonelli *et al*. 2015). This method ignores any other sources of rate variation and has been shown to be inadequate, especially for large phylogenies where substantial heterogeneity in diversification rates is expected (Caetano *et al*. 2018). One reason for the lack of attention to angiosperms is that their vast diversity makes it difficult to obtain well-sampled phylogenetic and distributional data at a species-level, which is required to estimate speciation rates accurately. Here we tested how speciation rates varied with latitudinal distribution in over 60,000 angiosperm species using different methods that accommodated macroevolutionary rate heterogeneity and incomplete taxon sampling. Our analyses used the largest phylogenetic trees presently available for angiosperms (Zanne *et al*. 2014; Smith & Brown 2018), alongside all geographic distribution data obtainable for these species (Mounce *et al*. 2017) from the Global Biodiversity Information Facility (GBIF). We show that latitudinal variation in recent speciation rates cannot generate the LDG because speciation was not faster in the tropics.

## Material and Methods

### Phylogenetic datasets and macroevolutionary rate estimates

We estimated speciation rates in the angiosperm radiation with Bayesian Analysis of Macroevolutionary Mixtures (BAMM) v.2.5.0 (Rabosky 2014). BAMM detects heterogeneity in speciation and extinction rates through time and across lineages. The input for BAMM was the most comprehensive time-calibrated phylogenetic tree for seed plants that was built using a hierarchical clustering analysis of Genbank sequence data (Smith & Brown 2018). On average, sister branches in this phylogeny had an overlap of 1792 base pairs, which corresponds to roughly one or two gene regions. We used a modified version of the tree containing GenBank taxa with a backbone provided by Open Tree of Life (i.e., GBOTB) for 72,986 flowering plants. Species names were standardised to The Plant List v.1.1 (Jin & Qian 2019). Following Igea *et al*. 2017, convergence of such a large BAMM analysis was achieved by dividing the initial tree into clades of less than 6000 species. Doing so resulted in 17 clades (16 monophyletic clades and 1 clade that contained the backbone of the tree plus the unassigned species), which we then used to run independent BAMM speciation/extinction analyses. Within each clade, non-random incomplete taxon sampling was incorporated into the estimation process by calculating the number of species sampled in each family. Prior settings were obtained using the *setBAMMpriors* function in the R package *BAMMtools* (Rabosky *et al*. 2014) and *expectedNumberOfShifts* was set at 50. Analyses were run for a minimum of 100 million generations and ≥20% of each run was discarded as a burn-in when necessary. Effective samples sizes of the log-likelihood and the number of rate shifts were both above 200. The event files for all 17 clades were then combined to generate a single BAMM result file that was used for all downstream analyses.

Although the reliability of BAMM estimates has recently been questioned (Moore *et al*. 2016), simulations have shown that accurate estimates of speciation can be obtained even in the absence of fossil data (Rabosky *et al*. 2017; Mitchell *et al*. 2018). Tip-based estimates of speciation like the ones we use here should also be robust to the effect of extinction and are increasingly used when analysing the effect of traits on lineage diversification and summarising geographic patterns of speciation rate variation (Title & Rabosky 2019). BAMM extinction estimates are thought to be more unreliable (Rabosky 2010; Marshall 2017; Mitchell *et al*. 2019), so we do not discuss them here. BAMM estimates of diversification rates, which are calculated as the net outcome of speciation minus extinction (Title & Rabosky 2019), will therefore also be more unreliable than speciation rates. Hence, and following other similar papers (Rabosky *et al*. 2018), we only discuss BAMM estimates of speciation. Furthermore, we also estimated recent speciation rates using the Diversification Rate metric (DR; Jetz *et al*. 2012, Supplementary Note 2), which only considers branch lengths and splitting events and does not accommodate for incomplete sampling. We also focus on estimates of speciation and extinction rates from other methods in subsequent sections.

### Latitudinal datasets

We obtained latitudinal data for 236,894 angiosperms from a study that estimated the median, maximum and minimum latitude for each species (Mounce *et al*. 2017) using observations from GBIF. Since our goal was to assess whether speciation rates drive the LDG, we focused on latitudinal differences and did not consider regional and longitudinal differences in diversity, which may be important in some cases (Reyes *et al*. 2015; Xing & Ree 2017; Igea & Tanentzap 2019). We then intersected these data with our BAMM dataset (n = 72,986) to obtain estimates of speciation rate and latitude for as many angiosperms as possible, hereafter referred to as the ‘full’ dataset (n = 60,990). The species names in the GBIF datasets were standardised with The Plant List (TPL) using the *Taxonstand* (Cayuela *et al*. 2012) and *Taxonlookup* (Pennell *et al*. 2016) packages before collation.

Tropical and temperate species were defined as those where the absolute median latitude was below and above 23.5°, respectively. Species were also grouped into equal-width latitudinal bins according to their median latitude to define latitudinal bands. To ensure that these latitudinal bands were completely tropical or temperate, the width of each band was set at 9.4° (i.e., starting from a band centered in the Equator from -4.7° to 4.7°, then from 4.7° to 14.1°, 14.1° to 23.5°, etc).

### BAMM-based correlation of tropicality and speciation

We correlated species-specific (i.e., tip-based) speciation rates from BAMM with the latitudinal data using Structured Rate Permutations on Phylogenies (STRAPP) (Rabosky & Huang 2016). STRAPP assesses the significance of the empirical association of macroevolutionary rates and phenotypic traits by comparing it to a null distribution generated by permuting the speciation rates across the phylogeny while maintaining the position of the rate shifts in the phylogenetic tree. Crucially, this test does not require that all rate variation is caused by the focal trait and thus shows a reduced Type I error rate (Rabosky & Huang 2016). We performed all STRAPP two-tailed tests using the *traitdependentBAMM* function of the *BAMMtools* package (Rabosky *et al*. 2014) with 1000 replicates and logging the rates. We assessed whether *i*) tropical and temperate species had different speciation rates using a Mann Whitney U test; *ii*) speciation rates for each of the latitudinal bands were different using the Kruskal-Wallis rank sum statistic; and *iii*) species absolute median latitude was correlated with speciation rate using Spearman’s rank correlation coefficient.

We repeated the analyses discarding sets of species that may bias our results. First, we used only densely sampled species (i.e., with five or more data points in the GBIF dataset). Second, we discarded widespread species (i.e., neither strictly tropical nor strictly temperate) for analyses comparing strict tropical and temperate taxa. Strictly tropical and temperate species were defined as species with median, maximum and minimum latitude all occurring either inside or outside the tropics, respectively. Third, we discarded species with median latitudes in the highest latitudinal bands (>50° and <-50°), which contained few species with extreme estimates of λ.

### Assessing the reliability of the BAMM results

We assessed the consistency of the BAMM results in the face of geographic biases, low sampling and topological uncertainty.

First, the full dataset had more temperate than tropical species and so was inconsistent with the LDG. Poorly sampled clades may have average longer branches and lower λ and so our speciation rate estimates may be affected by latitudinal sampling bias. We therefore tested whether clades with lower sampling fractions were found at lower latitudes and had lower λ rates. We employed two additional strategies to analyse the effect of the disproportionate amount of temperate species in our BAMM analyses. We generated 100 random samples of our full dataset (herein ‘unbiased’ datasets), each with 30,000 species that maintained the proportion of tropical (57.5%) and temperate (42.5%) species present in GBIF. We also generated 100 random samples (herein ‘extreme tropical’ datasets) that assumed all species absent from the GBIF dataset of Mounce *et al*. (2017) were tropical, so the proportions of tropical and temperate species were 70.9% and 29.1%, respectively. We derived these values by estimating that 109,461 species were absent from GBIF by subtracting the number of species in the GBIF dataset from the number of species in The Plant List according to *TaxonLookup* (Pennell *et al*. 2016). Although both the unbiased and extreme tropical datasets were generated to assess the effect of excess temperate species, the extreme tropical dataset also represented the most extreme scenario of undescribed biodiversity. The sizes of these datasets was fixed at 30,000 species so that the tropical species present in each subsample were not excessively redundant when obtained from a total pool of 26,873 species by random draws of 17,130 and 21,273 species for the unbiased and extreme tropical datasets, respectively. Using the two datasets, we then assessed whether the full dataset λ estimates were affected by poorer sampling for tropical species. We did so by selecting 10 random replicates of the 30,000-species subsampled datasets and rerunning the BAMM speciation analyses as detailed above. We determined whether the relationship of the full with both the unbiased and extreme tropical λ estimates was different for tropical and temperate species. To do this, we predicted the full λ estimates using the subsampled λ estimates, ‘tropicality’ (i.e. a binary variable indicating if a species was exclusively tropical based on our aforementioned definition) and the interaction of the λ estimates and tropicality. We fitted this model separately for each of the 10 replicates in the unbiased and extreme tropical datasets, incorporating the effect of the phylogeny using the *phylolm* package (Tung Ho & Ané 2014). We also tested whether the association between speciation and latitude in the full dataset stemmed from the excessive proportion of temperate species by repeating the STRAPP analyses with the 100 replicates in the unbiased and extreme tropical datasets.

Second, our full dataset contained λ estimates for 60,990 species, which represent 17.6% of described angiosperm species (total n = 346,365 calculated with *Taxonlookup*). To assess the reliability of λ estimates given the large number of missing taxa, we randomly generated 10 subsamples (‘small’ datasets) with the same proportion (i.e., 17.6%) of the species in our full dataset (n = 10,739). As above, we ran BAMM for each of the 10 subsamples and predicted the full λ estimates using the small λ estimates.

Third, we analysed a single phylogenetic tree, which may only represent one of many potential hypotheses for the angiosperm ToL. To assess the effect of the unaccounted topological and branch length uncertainty, we tested the relationship between speciation rates and latitude in a smaller dataset of 28,057 species obtained from an independently estimated phylogenetic tree. The tree was estimated with GenBank sequences for seven gene regions with a median per-taxon proportion of missing data of 0.60 (Zanne *et al*. 2014; Qian & Jin 2016). We obtained BAMM λ rates for this dataset as described elsewhere (Igea *et al*. 2017). We then tested the correlation between the BAMM estimates of our full dataset and this reduced dataset. We also used the reduced dataset to determine whether there were differences in BAMM λ rates between tropical and temperate species and among different latitudinal bands; and whether absolute median latitude and speciation rates were associated. All of these analyses were performed as described above.

### Geography-dependent speciation

We used the Geographic Hidden-State Speciation and Extinction (GeoHiSSE) framework to test for further differences in speciation between tropical and temperate lineages. GeoHiSSE estimates speciation, extinction and transition rates for two geographic areas while incorporating rate heterogeneity that is independent of geography by including unobserved characters known as “hidden states” (Caetano *et al*. 2018). Including these hidden states is crucial to create biologically meaningful null models where diversification varies independently of the focal traits. We considered any species occurring strictly below and above 23.5° as tropical and temperate respectively, while species whose range span both >23.5° and <23.5° were classified as widespread. Sampling fractions for each geographic state in the phylogenetic tree were estimated to be 0.12, 0.30, and 0.24 for strictly tropical, strictly temperate, and widespread species, respectively. Following Caetano *et al*. 2018, we fitted 12 different models (numbers 1-12 in Caetano *et al*. 2018, Table 1). Some of these were “null models” where no diversification rate variation is caused by geography. Other models included area-dependent and area-independent diversification with two to five hidden states, and some also separated range contraction from lineage extinction (i.e., “+extirpation models”). We then selected the best fitting model using the sample-sized corrected Akaike Information Criterion (AICc). We used the “fast version” of the GeoHiSSE model as implemented in *hisse* v.1.9.6 (Beaulieu & O’Meara 2016) to make analyses feasible in such a large phylogeny. The size of the dataset meant that some of the models took a very long time to fit (2 months on 2.5 GHz processor for a single fit). We could therefore not obtain rate estimates across the nodes and tips using the marginal reconstruction algorithm, which required estimating probabilities for all possible combinations of states at each node in the tree given all combinations at nodes in the rest of the tree. It was also not feasible to model-average the “effective state proportion” (i.e., the estimated proportion of each geographic area and hidden state combination), which would have allowed us to test if a only a small proportion of the lineages were associated with a geographic-dependent diversification process (Caetano *et al*. 2018).

**Table 1.** Model comparison of 12 GeoHiSSE models with up to five hidden rate categories. Model numbers and descriptions refer to Caetano *et al* (2018). Null models have no differences in diversification associated with geography and diversification parameters are constrained to be equal between areas in the same hidden state category. Area independent models (CID) have no geographic-dependent differences in diversification. Full models have no constrained parameters. Extremely high AICc values for the three simplest models suggest that they were a very poor fit. Original GeoSSE refers to the formulation in Goldberg *et al*. (2011)

| model | description | # free parameters | AICc |
| --- | --- | --- | --- |
| 5 | CID-GeoHiSSE, 5 hidden rate classes, null model | 13 | 418090 |
| 11 | CID-GeoHiSSE+extirpation, 5 hidden rate classes, null model | 15 | 423888 |
| 10 | GeoHiSSE+extirpation, 2 hidden rate classes, full model | 19 | 425927 |
| 9 | CID-GeoHiSSE+extirpation, 3 hidden rate classes, null model | 11 | 428085 |
| 12 | CID-GeoHiSSE+extirpation, 2 hidden rate classes | 9 | 429549 |
| 4 | GeoHiSSE, 2 rate classes, full model | 15 | 430692 |
| 3 | CID-GeoHiSSE, 3 hidden rate classes, null model | 9 | 442036 |
| 8 | GeoSSE+extirpation, full model | 9 | 458691 |
| 2 | Original GeoSSE, full model | 7 | 470420 |
| 1 | CID-original GeoSSE | 4 | 1.56E+19 |
| 6 | CID-GeoHiSSE, 2 hidden rate classes | 7 | 1.56E+19 |
| 7 | CID-GeoSSE+extirpation | 6 | 1.56E+19 |

### Clade-based analyses

We estimated clade-level measures of speciation across the angiosperm phylogenetic tree. We used a set of 4 million-year-wide time intervals from 0 to 24 million years (myr) to define the ages of the clades. Clades were delimited as the largest non-overlapping monophyletic groups of four or more species where sampling was larger than 30%. This criterion ensures that the probability of retrieving the correct crown age of the clade is above 70% (Rabosky 2016). At least 50% of the species in the clades also had to have latitudinal data. We used RPANDA (R: Phylogenetic ANalyses of DiversificAtion, Morlon *et al*. 2016) to fit six different models of diversification: *i*) constant speciation (λ) and extinction (μ) fixed at 0; *ii*) constant λ and μ; *iii*) exponential λ and μ fixed at 0; *iv*) exponential λ and constant μ; *v*) constant λ and exponential μ; and *vi*) exponential λ and μ. Incomplete sampling within clades was incorporated into the estimates by calculating the proportion of sampled species within the genera in each clade. Speciation rates were estimated using model averaging with AIC weights. The correlation of the clade-level estimates of speciation and the proportion of temperate and tropical species in each clade was assessed using phylogenetic least squares regression as implemented in the R package *nlme* (Pinheiro *et al*. 2014). The lambda parameter was optimised using a maximum likelihood approach. We weighted the regressions by the proportion of GBIF data points in each clade to account for different sampling levels. We also assessed if potential latitudinal differences in sampling proportions could explain our results. We did so by testing if the sampling proportions of largely tropical and largely temperate clades (defined as clades where >50% of the species were tropical and temperate, respectively) were different across the 6 time intervals using Pearson’s chi-squared test.

## Results

We found that tip-specific speciation rates were consistently smaller in the tropics for 60,990 species across the whole angiosperm radiation. These results were repeatable with several ways of classifying species as tropical or temperate. Using correlation tests that controlled for phylogenetic pseudoreplication (Rabosky & Huang 2016), we found that tropical species had smaller mean speciation rates (λ) than temperate species (Fig 1a; λ_temperate_ = 0.73 species/myr, λ_tropical_ = 0.65 species/myr, p-value = 0.043). Angiosperms in latitudinal bands closer to the poles (i.e. absolute median latitude >42.3°) speciated at faster rates than in any tropical latitudinal band (Fig. 1b). The absolute median latitude occupied by individual species was also positively correlated with their speciation rate (Fig. 1c; Spearman’s ρ = 0.086, p-value = 0.009). These results were maintained when we: *i*) analysed only those species that were densely sampled in GBIF (n = 53,344 species) and *ii*) discarded the highest latitudinal bands (>50° N and S), which contained fewer species (n = 60,034) (Table S1).

**Figure 1.**
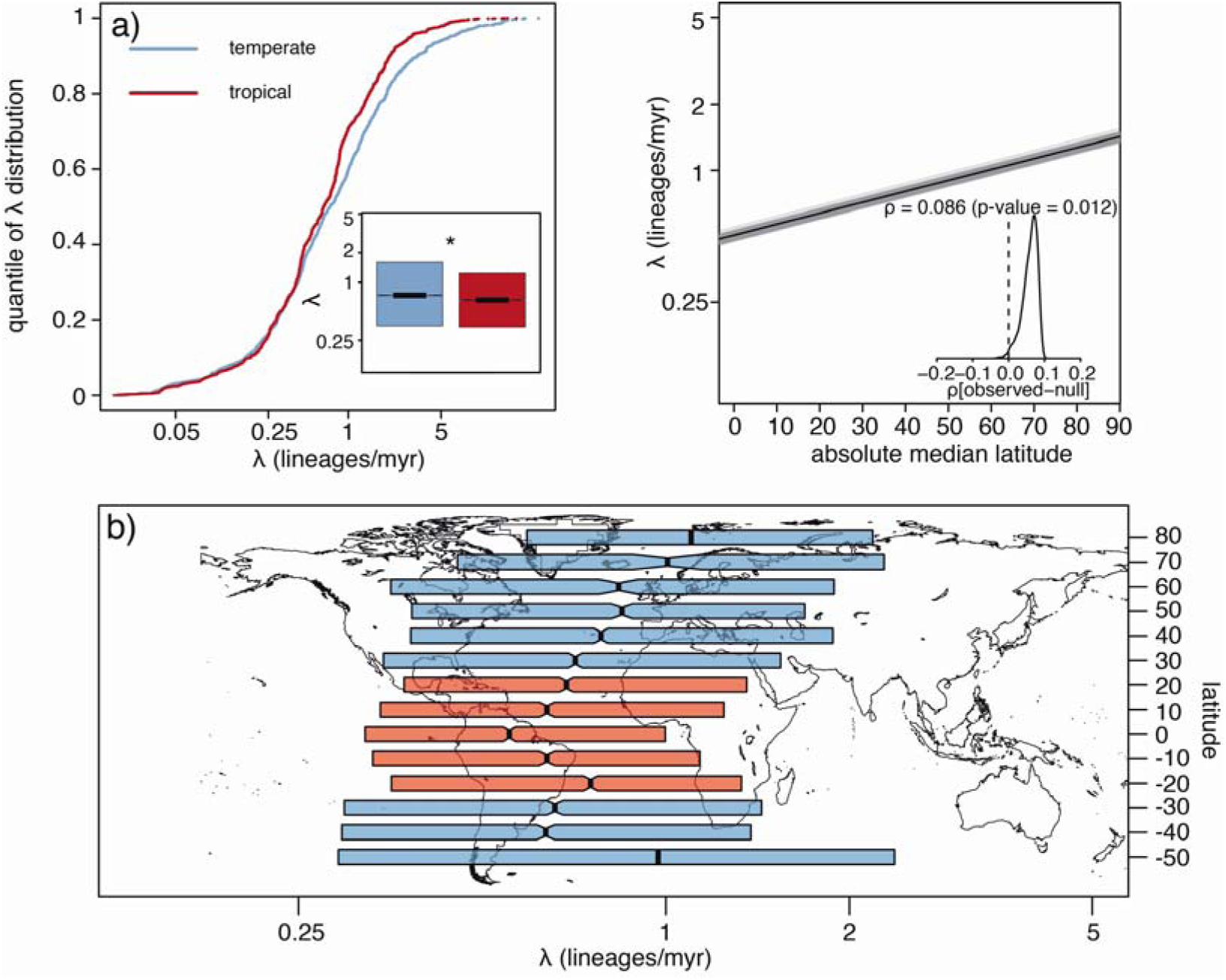
Tropical species have smaller speciation rates (λ) than temperate species. **a)** Rank-ordered distribution and boxplot (inset) of tip λ inferred with BAMM for tropical (red) and temperate (blue) species. * indicates significant difference (p-value = 0.043) between the two groups, assessed with a Mann-Whitney test implemented in STRAPP. **b)** λ grouped by latitudinal band of each species. Notches in boxplots indicate 95% confidence intervals around median, denoted by thick vertical lines and boxes span the interquartile ranges. **c)** Spearman’s ρ correlation between species absolute median latitude and λ as estimated with STRAPP. Grey lines are correlations across 1000 samples of the posterior distribution estimated with BAMM. Solid black line indicates median correlation ± 95% confidence interval across the posterior distribution. Inset shows the difference between the empirical and null correlations estimated with 1000 permutations of evolutionary rates across the phylogeny.

Speciation rates were also different when comparing strictly tropical and strictly temperate species by discarding species that occurred both inside and outside the tropics (n = 46,426), but this difference was not significant (p-value = 0.064; Table S1). Our full dataset contained a disproportionate amount of temperate species and so did not hold a LDG (Fig. S1), but this bias could not explain the consistent association between latitude and speciation rates. First, we generated 100 “unbiased” datasets that maintained the proportion of tropical and temperate species present in GBIF and were consistent with the LDG. Second, we generated another set of 100 “extreme tropical” datasets, where the proportions of temperate and tropical species were 29.1% and 70.9%, respectively. Rerunning BAMM for 10 of the 100 unbiased and extreme tropical datasets, we found that the λ estimates were very positively correlated with those from the full dataset (median ρ = 0.82, median p-value < 0.001 for both the unbiased and the extreme tropical datasets; Figs 2, S2). This correlation between λ of the full and subsampled datasets did not differ between tropical and temperate species, as the effect size of the interaction between tropicality and the subsampled λ estimates was not different from zero across all the 10 replicates for the unbiased (mean = -0.0005, standard deviation = 0.0017) and the extreme tropical (mean = -0.0015, standard deviation = 0.0024) datasets. As with the full dataset, we also found that speciation rates in the tropics were consistently lower and that λ was positively correlated with absolute median latitude using STRAPP across the 100 replicates of both the unbiased (Fig. 3) and extreme tropical (Fig. S3) datasets. Finally, we confirmed that our results could not be explained by better sampled temperate species having higher λ. Densely sampled clades were only slightly more frequent at higher latitudes (Fig. S4), and they had smaller estimates of λ (Fig. S5), contrary to what we might expect if our results were a sampling artefact.

**Figure 2.**
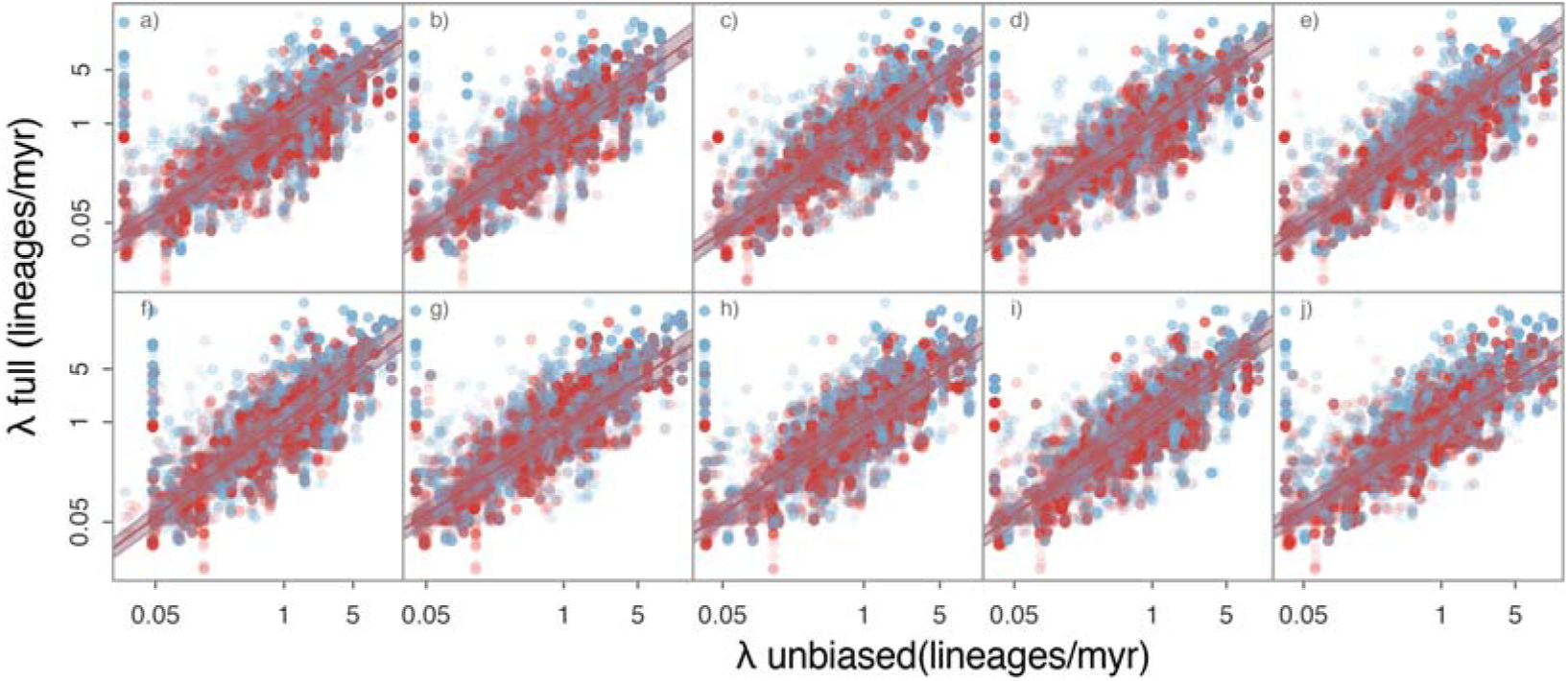
BAMM λ estimates were positively correlated in the full and 10 unbiased datasets (a to j) with no effect of tropicality. Solid lines are phylogenetic linear regressions predicting λ in the full tree (n = 60,990 species) with λ in the unbiased tree (n = 30,000 species) in temperate (shown in blue) and tropical (shown in red) species. Shaded areas indicate the 95% confidence intervals.

**Figure 3.**
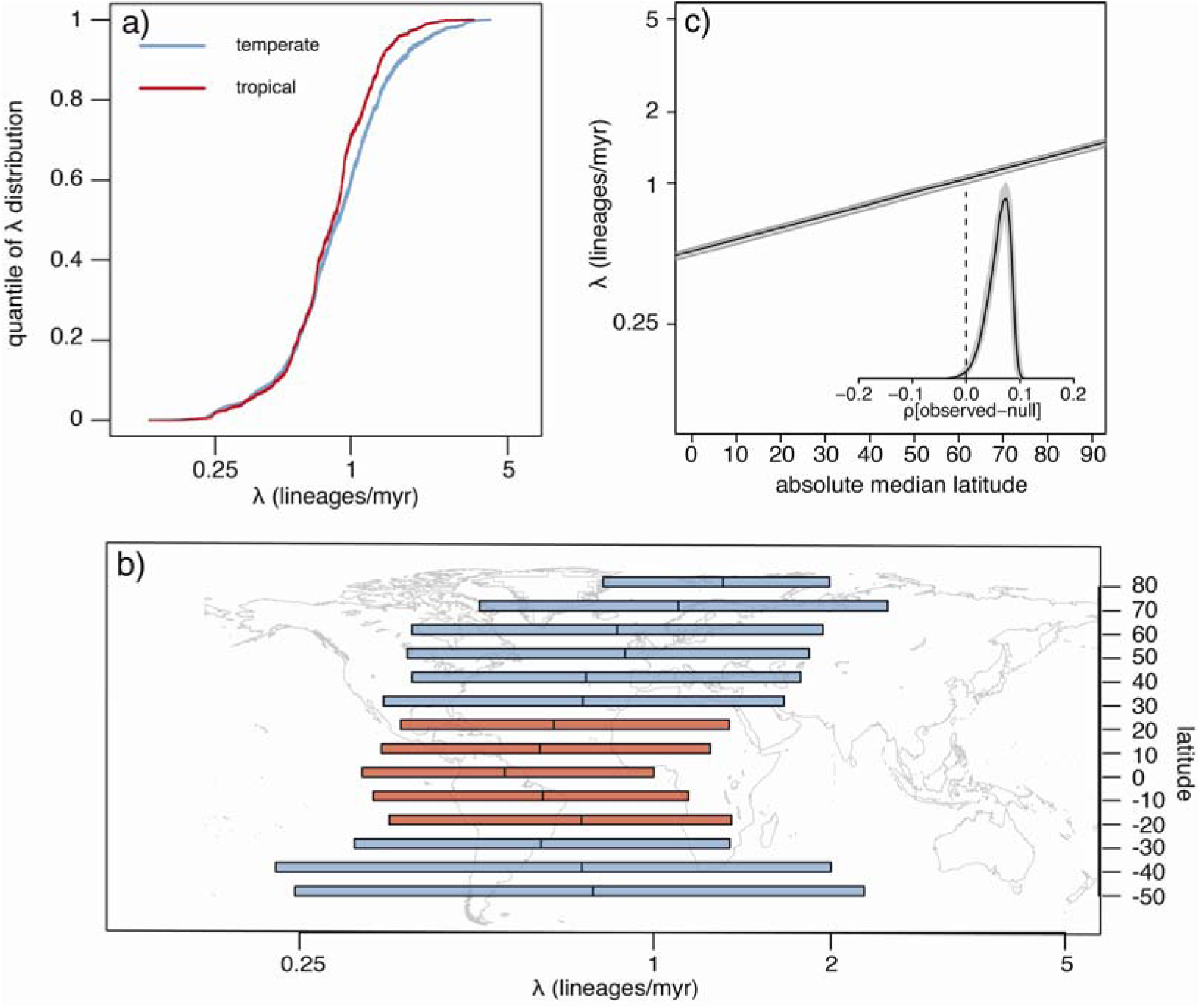
Temperate species have higher speciation rates (λ) than tropical species for 100 subsampled “unbiased” datasets. Datasets were generated whereby 42.5% and 57.5% of species were temperate and tropical, respectively. **a)** Rank-ordered distribution of λ inferred with BAMM for tropical (red) and temperate (blue) species. **b)** λ grouped by latitudinal band of each species. Boxes show the average interquartile range across the 100 subsampled datasets, and lines show the average medians. **c)** Spearman’s ρ correlation of species absolute median latitude and λ as estimated with STRAPP. Solid black line indicates median correlation ± 95% confidence interval across the posterior distribution. Inset shows the median (black line) ± 95% confidence interval for the difference between the empirical and null correlations estimated with 1000 permutations of evolutionary rates across the phylogeny in each of 100 subsampled datasets.

We also confirmed that the speciation rate estimates were robust to the low levels of taxon sampling. We compared the BAMM λ estimates from the full dataset with estimates generated by randomly sampling 17.6% of the species in our full dataset 10 times and rerunning BAMM. This level of sampling reflected the proportion of angiosperm species included in our full dataset from those described in The Plant List. The resulting λ estimates from the full and the small datasets were positively correlated (median ρ = 0.67, median p-value < 0.001). This association was slightly weaker for tropical than for temperate species, but this difference was very small and not statistically significant across the 10 replicates (mean = -0.04, standard deviation = 0.03; Fig. S6).

Our results were based on a single phylogenetic tree that represented one of many possible topologies in the angiosperm ToL, and so we confirmed that the results were consistent with another independently derived phylogeny (Zanne *et al*. 2014; Qian & Jin 2016). Using BAMM λ estimates and latitudinal information for this 28,057 species tree, we found that λ was higher outside the tropics and that speciation rates increased towards the poles (Fig. S7). The BAMM λ estimates of this topology and our full dataset were positively correlated (Spearman’s ρ = 0.46, p-value < 0.001).

Elevated speciation rates in the tropics were also rejected as the driving force of the LDG using a method that explicitly tested the influence of geography on species diversification. The best fitting model with GeoHiSSE (AICc weight = 1; Table 1) had five different hidden states that accommodated rate variation across lineages and no differences in turnover and extinction fraction between tropical and temperate regions. This result suggests that most of the variation in diversification rates across the angiosperm radiation is not linked to latitude, though some angiosperm lineages can show range-dependent diversification. We were precluded from exploring this latter process using marginal reconstructions by the sheer size of the dataset (see Methods). The confidence intervals around the maximum-likelihood estimates were too wide to identify any differences in rate estimates, so we do not discuss them further.

Finally, we found that tropical species did not speciate faster across clades of different ages in the angiosperm phylogeny. Using clades in six 4 million-year-wide time slices, from 0 to 24 million years, where we were confident in speciation rates and GBIF coverage, we fitted models that implemented time-variable and time-constant speciation and extinction. Mirroring the BAMM results, clades with a larger proportion of temperate species had higher rates of speciation in the best fitting model (Fig S8). This result was not totally explained by sampling biases. Temperate clades had significantly denser average sampling only in the 8-12 and 12-16 myr intervals (χ^2^ = 5.22, df = 1, p-value = 0.020 and χ^2^ = 12.19, df = 1 p-value < 0.001, respectively), and these differences were small (0.65 vs 0.62 and 0.65 vs 0.61 for temperate vs tropical in each period, respectively).

## Discussion

Using data for over 60,000 flowering plants, our findings rejected the long-standing notion that the greater species diversity of the tropics can be explained by higher recent speciation rates (Mittelbach *et al*. 2007), such as from higher environmental energy (Dowle *et al*. 2013). We instead found higher rates of recent speciation closer to the poles. An explanation for this difference is that temperate biotas have fewer species, so their niche space may be less saturated, thereby increasing opportunities for lineage divergence (Simpson 1953; Schluter 2016). Similarly, reproductive isolation may be elevated at higher latitudes by the greater ecological opportunity generated from recurrent environmental change and climate instability (Cutter & Gray 2016). Other explanations may be related to the ecological and life history traits of temperate species. For example, faster speciation rates at higher latitudes could stem from the higher frequency of small-seeded species (Moles *et al*. 2007). Small seed size is positively correlated with angiosperm diversification (Igea *et al*. 2017), and temperate species did have smaller seeds in our full dataset (Supplementary Note 1). Therefore, our results suggest that latitudinal differences in recent speciation rates do not shape the LDG but are shaped by differences in species diversity and traits (Weir & Price 2011; Schluter & Pennell 2017).

Rejecting the role of speciation rate as the cause of the LDG calls for alternative explanations (Antonelli *et al*. 2015). Many theories have been proposed to explain the LDG, but they can all be divided into four broad categories arising from differences in: *i)* diversification rates; *ii*) dispersal rates; *iii)* time for species accumulation and *iv)* ecological limits to species diversity (Etienne *et al*. 2019). Hypotheses in the different categories often invoke multiple causal mechanisms and so may not be mutually exclusive (Pontarp *et al*. 2019). First, the elevated speciation near the poles could be coupled with higher extinction rates to produce a scenario where temperate zones act as both a “cradle” and “grave” of biodiversity (Cutter & Gray 2016). Net diversification might therefore still be higher in tropical areas despite the absolute rates of speciation and extinction each being smaller than closer to the poles. Our GeoHiSSE analyses do not, however, support this scenario as we found no such differences in turnover (Table 1). Estimates of extinction rate from molecular phylogenies in the absence of fossil data are often unreliable (Marshall 2017; Mitchell *et al*. 2018), which further complicates direct tests of this hypothesis. Second, the tropical conservatism hypothesis proposes that lower net migration from the tropics to temperate zones contributes to the LDG (Wiens & Donoghue 2004), because most lineages originated in the tropics and have been unable to disperse into harsher, temperate environments. For example, in New World woody angiosperms, transitions between tropical and temperate environments are rare and most temperate lineages are relatively young (Kerkhoff *et al*. 2014). This scenario was also not supported with GeoHiSSE as we detected no differences in transition rates between the tropics and the temperate zones and vice versa. Third, if rates of diversification and migration have no latitudinal variation, the greater diversity of the tropics may stem from their larger area through time, i.e. the age-and-area hypothesis (Fine & Ree 2006). For instance, current species richness across the world’s 32 bioregions is best predicted by area and productivity over geological time (Jetz & Fine 2012). Area-through-time differences have also been proposed to underlie differences in diversity among tropical regions (Couvreur 2015). Fourth, a whole class of hypotheses posits that there are ecological limits to the number of species that a region can hold, and that these limits vary with latitude (Rabosky 2009). For example, the larger diversity in the tropics could be explained by larger niche spaces and greater niche packing (Pellissier *et al*. 2018). Such a mechanism could interact with differential diversification rates to amplify spatial differences in diversity (Etienne et al. 2019). As ecological and macroevolutionary processes can interact, integrating all of them into a mechanistic eco-evolutionary framework can ultimately advance the search for explanations to the LDG (Pontarp *et al*. 2019). Other ecological interactions, like negative conspecific density dependence arising from herbivores and pathogens, have also been proposed to explain the high diversity in the tropics (LaManna *et al*. 2017).

Recent findings have suggested that rates of species origination are highest in species-poor areas (Quintero & Jetz 2018; Rabosky *et al*. 2018). Our results are consistent with these findings and therefore suggest that latitudinal variation in recent speciation rates is not a major engine of global diversity patterns in one of the largest eukaryotic radiations. Using time-heterogeneous estimates of speciation, we were able to exclude faster rates of evolution, such as arising from more favourable environmental conditions or more intense biotic interactions, as responsible for shaping the angiosperm LDG. Future work must now resolve whether recent rapid speciation nearer the poles is widespread across the ToL and its underlying causes.

## SUPPLEMENTARY MATERIAL

**Table S1.** Temperate species have mean higher speciation rates (λ, estimated with STRAPP) than tropical species across different datasets: “more.GBIF.data” excludes species with less than five GBIF data points; “no.widespread” excludes species occurring both in and outside the tropics; “no.high.latitude” excludes species from poorly sampled latitudinal bands with absolute median latitude ≥50°. Units for λ and dataset size are lineages/myr and species, respectively.

| dataset name | $\lambda_{\text{temperate}}$ | $\lambda_{\text{tropical}}$ | p-value | dataset size |
| --- | --- | --- | --- | --- |
| more.GBIF.data | 0.702 | 0.641 | 0.050 | 53,344 |
| no.widespread | 0.760 | 0.668 | 0.064 | 46,426 |
| no.high.latitude | 0.719 | 0.652 | 0.044 | 60,034 |

### Supplementary Note 1

Smaller seeded species have been shown to speciate faster than large seeded species (Igea *et al*. 2017). We assessed whether seed size differed between temperate and tropical lineages as it has previously been shown to do so (Moles *et al*. 2007), and so latitudinal variation in seed size could explain the differences in speciation between tropical and temperate species. We obtained seed size measurements for 13,178 species in our full dataset from a previous study (Igea *et al*. 2017) and we found that seed size was larger in tropical species (mean seed size_temperate_ = 0.0018 g; mean seed size_tropical_ = 0.0125 g; t = -28.239; df = 4300.7; p-value < 0.001). This difference remained when phylogeny was considered (phylANOVA: t = 33.933; p value = 0.001; significance assessed with 1,000 random simulations using phytools (Revell 2012))

### Supplementary Note 2

BAMM is a model-based approach to estimate tip speciation rates. To compare the BAMM results with a non-model based approach (i.e., that only relies on branch lengths and splitting events), we calculated the Diversification Rate metric (DR; Jetz *et al*. 2012) for the species in our full dataset (n = 60,990). This measure of recent speciation rate incorporates the number of nodes and the internode distances separating a species from the root and gives greater weight to branches closer to the present (Title & Rabosky 2019). We then compared the DR values of tropical and temperate species and assessed the variation of DR across latitudinal bands. Mirroring the BAMM results, we found that tropical species had smaller values of DR and that latitudinal bands closer to the poles had higher values of DR (Fig. S9).

**Figure S1.**
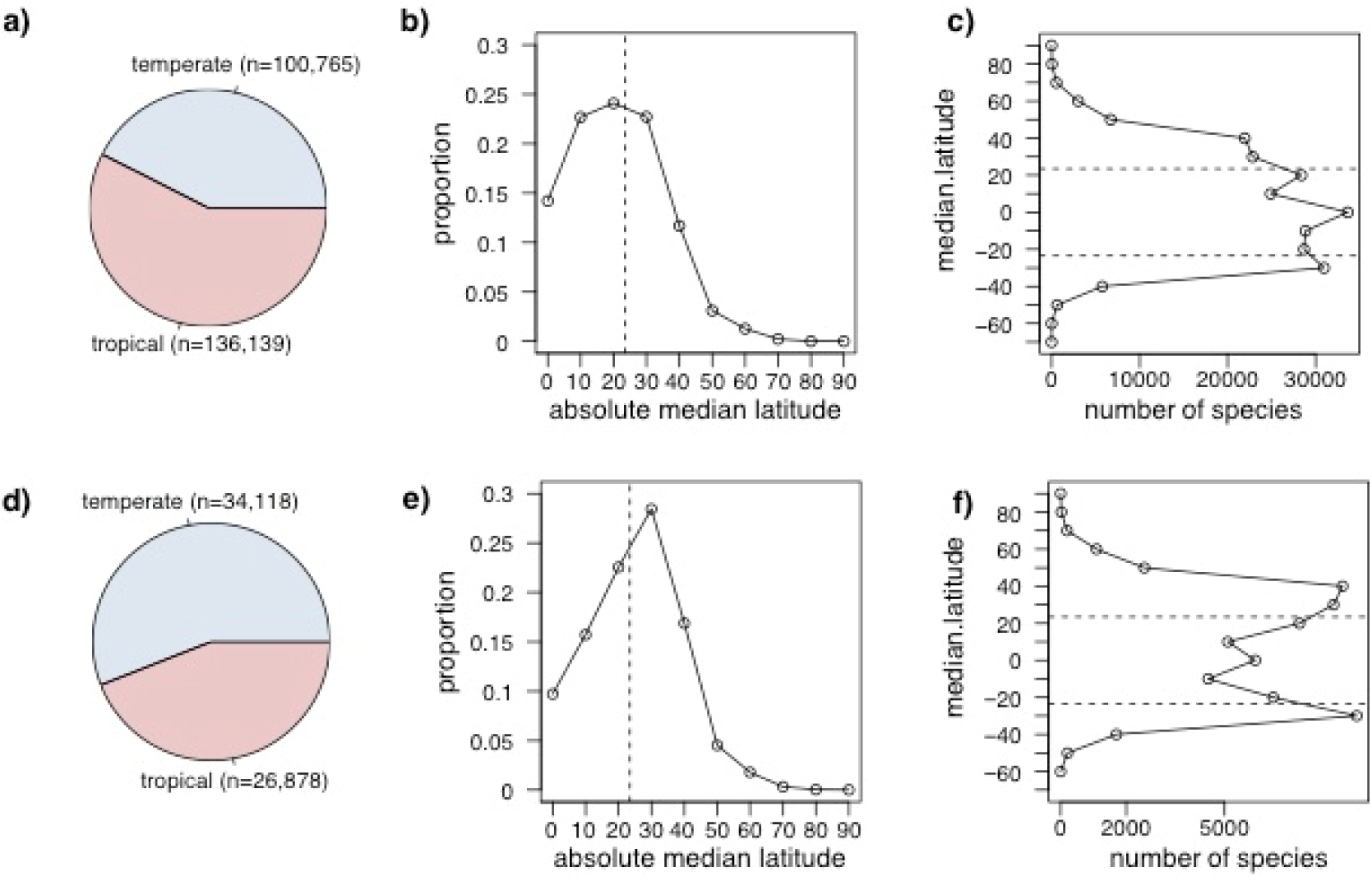
The full dataset had a disproportionate number of temperate species. **a)** Proportion of tropical and temperate species in the GBIF dataset (n = 236,894); **b)** proportion of species in each latitudinal band in the GBIF dataset; **c)** number of species in each latitudinal band in the GBIF dataset; **d)** proportion of tropical and temperate species in the full dataset (n = 60,990); **e)** proportion of species in each latitudinal band in the full dataset; **f)** number of species in each latitudinal band in the full dataset; in **b), c), e)** and **f)**, the dotted vertical line denotes the 23.5° threshold between tropical and temperate zones.

**Figure 2.**
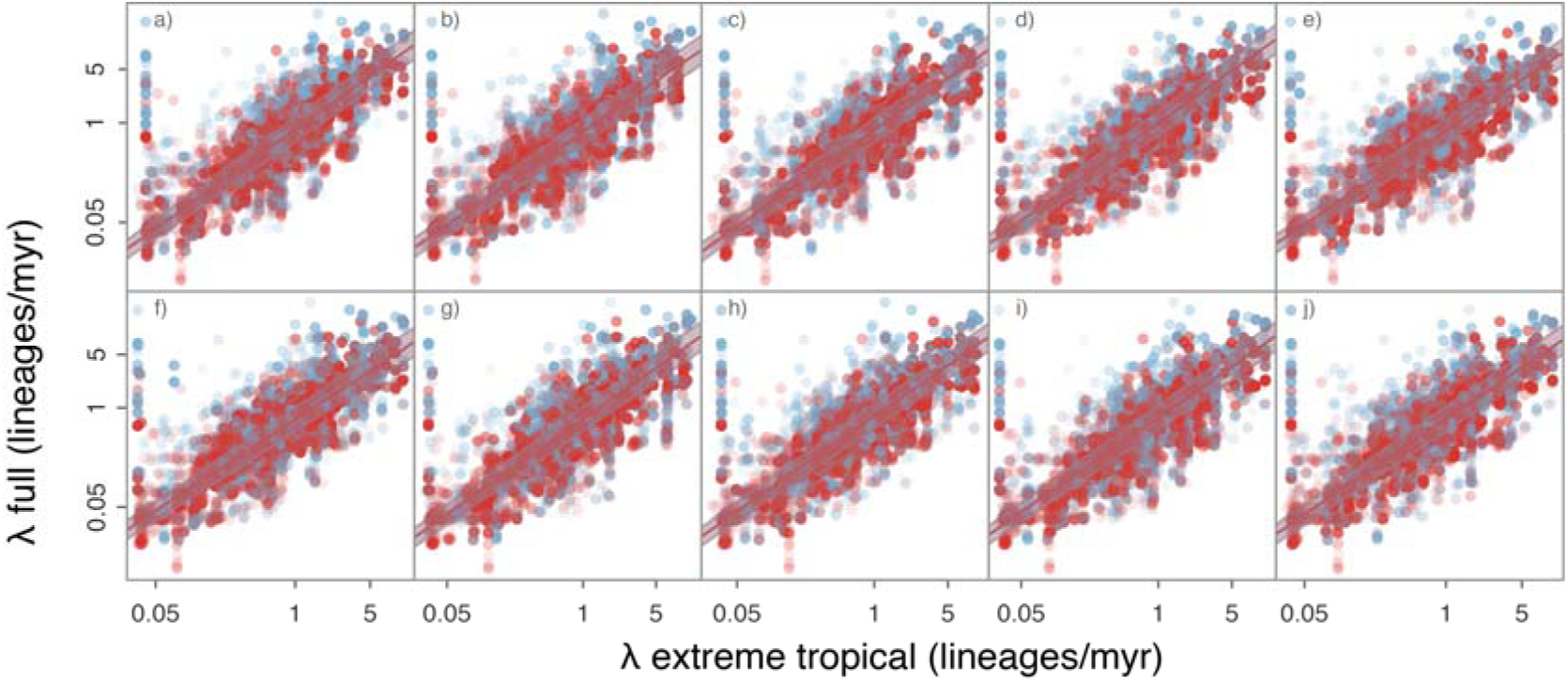
BAMM λ estimates were positively correlated in the full and 10 extreme tropical datasets (a to j) with no effect of tropicality on this correlation. Solid lines are phylogenetic linear regressions predicting λ in the full tree (n = 60,990 species) with λ in the extreme tropical tree (n = 30,000 species) in temperate (blue) and tropical (red) species. Shaded areas indicate the 95% confidence intervals.

**Figure S3.**
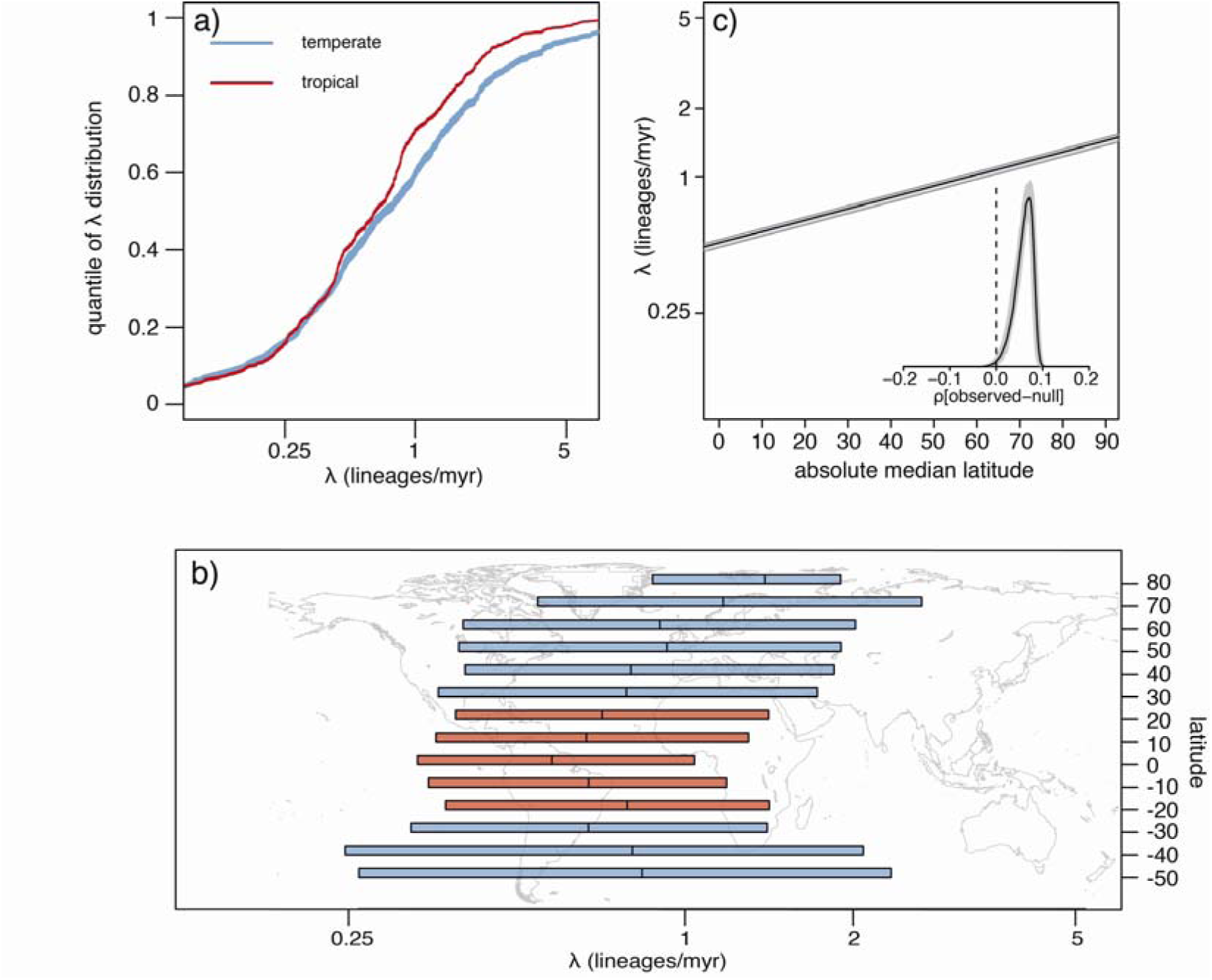
Temperate species have higher speciation rates (λ) than tropical species for 100 subsampled “extreme tropical” datasets with 29.1% temperate and 70.9% tropical species. **a)** Rank ordered distribution of λ inferred with BAMM for tropical (red) and temperate (blue) species. **b)** λ grouped by latitudinal band of each species. Boxes and lines are presented as in Fig. S4. **c)** Spearman’s ρ correlation of species absolute median latitude and λ as estimated with STRAPP. . Solid black line indicates median correlation ± 95% confidence interval across the posterior distribution. The inset shows the median (black line) ± 95% confidence interval for the difference between the empirical and null correlations estimated with 1000 permutations of evolutionary rates across the phylogeny in each of 100 subsampled datasets.

**Figure S4.**
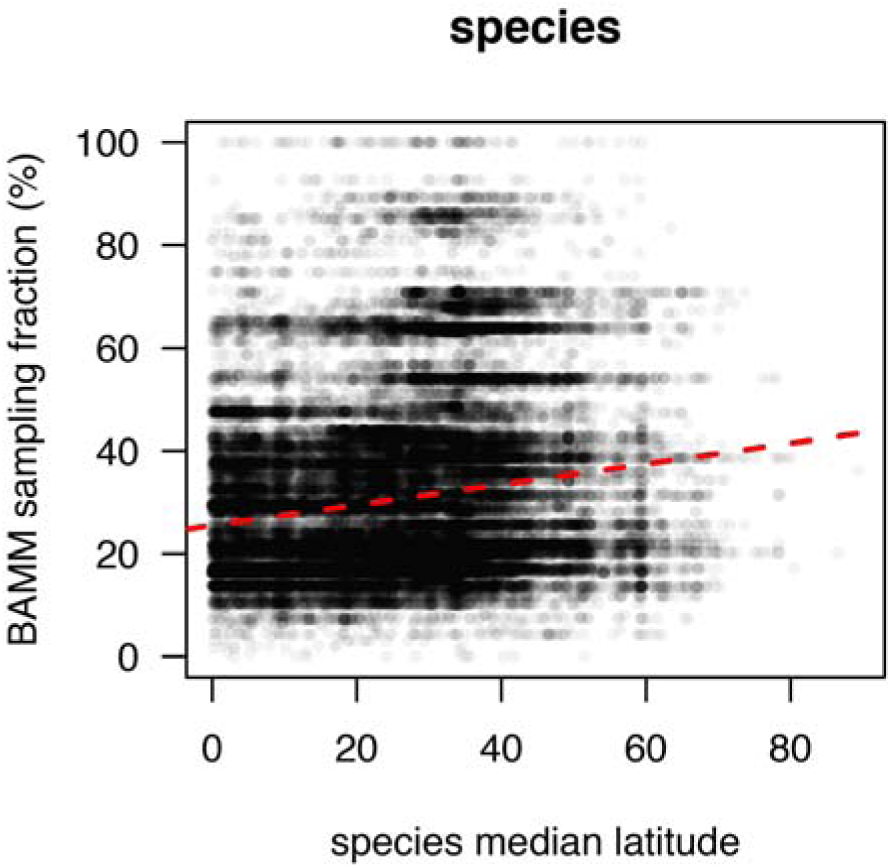
Relationship between the species absolute median latitude and the family-level sampling fraction used in the BAMM analyses. The red line is the slope of the linear regression (slope = 0.201, p-value < 0.0001).

**Figure S5.**
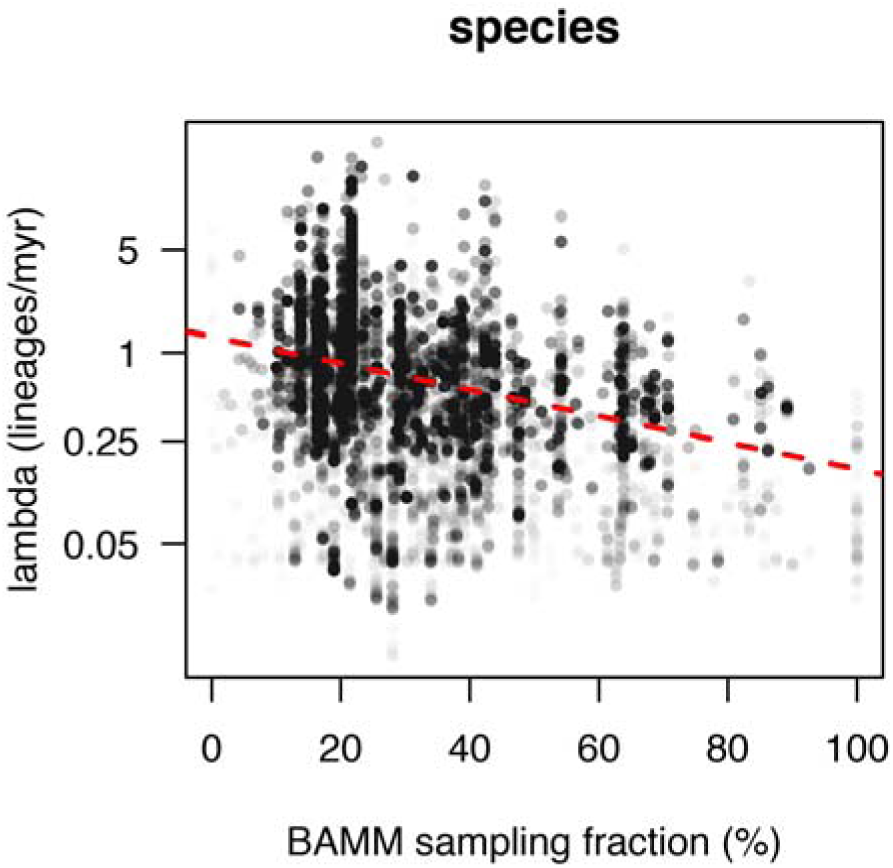
More densely sampled clades have smaller estimates of speciation rate (λ). The dotted line is the slope of the linear regression of the log(λ) and the BAMM sampling fraction (slope = -0.027, p-value < 0.0001).

**Figure S6.**
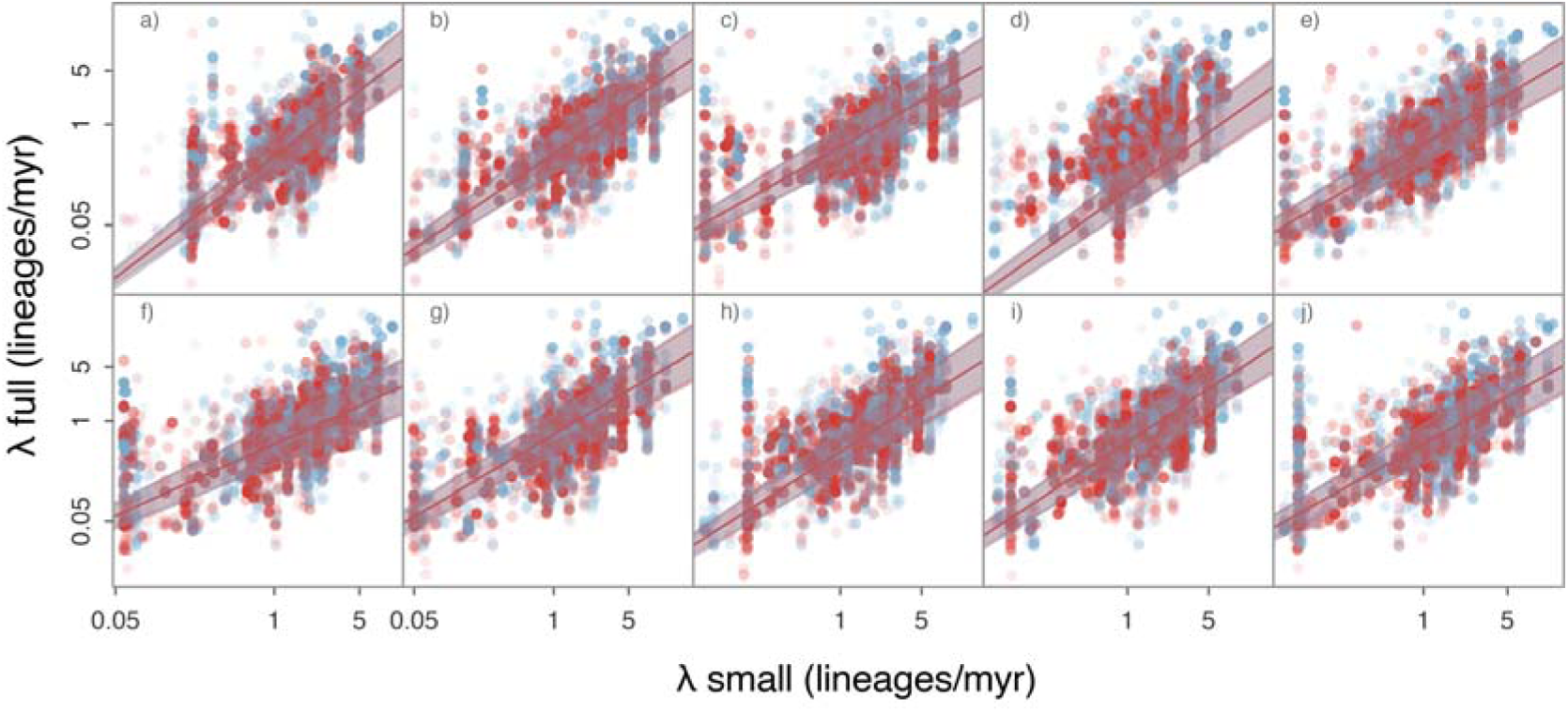
BAMM λ estimates are positively correlated in the full and 10 small datasets and tropicality has no effect on this correlation. **a) - j)** Solid lines are phylogenetic linear regressions predicting λ in the full tree (n = 60,990 species) with λ in the small tree (n = 10,739 species) in temperate (shown in blue) and tropical (shown in red) species. Shaded areas indicate the 95% confidence intervals.

**Figure S7.**
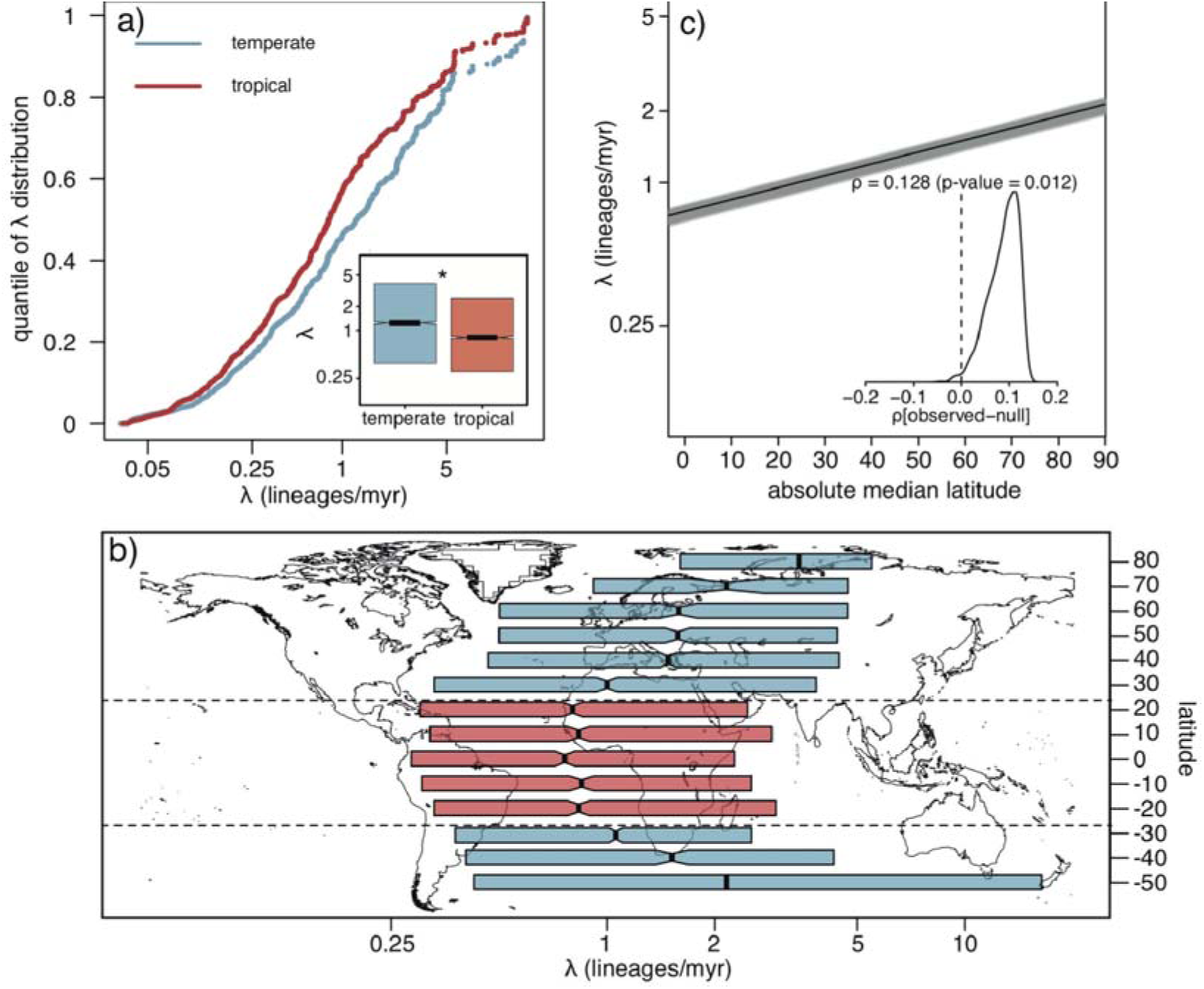
Tropical species have smaller speciation rates (λ) than temperate species with the Zanne phylogeny (n = 28,057 species). a) Rank ordered distribution and boxplot (inset) of λ inferred with BAMM for tropical (red) and temperate (blue) species; b) λ grouped by latitudinal band of each species; and c) Spearman’s ρ correlation between species absolute median latitude and λ as estimated with STRAPP. Lines, boxes and symbols as in Fig. 1.

**Figure S8.**
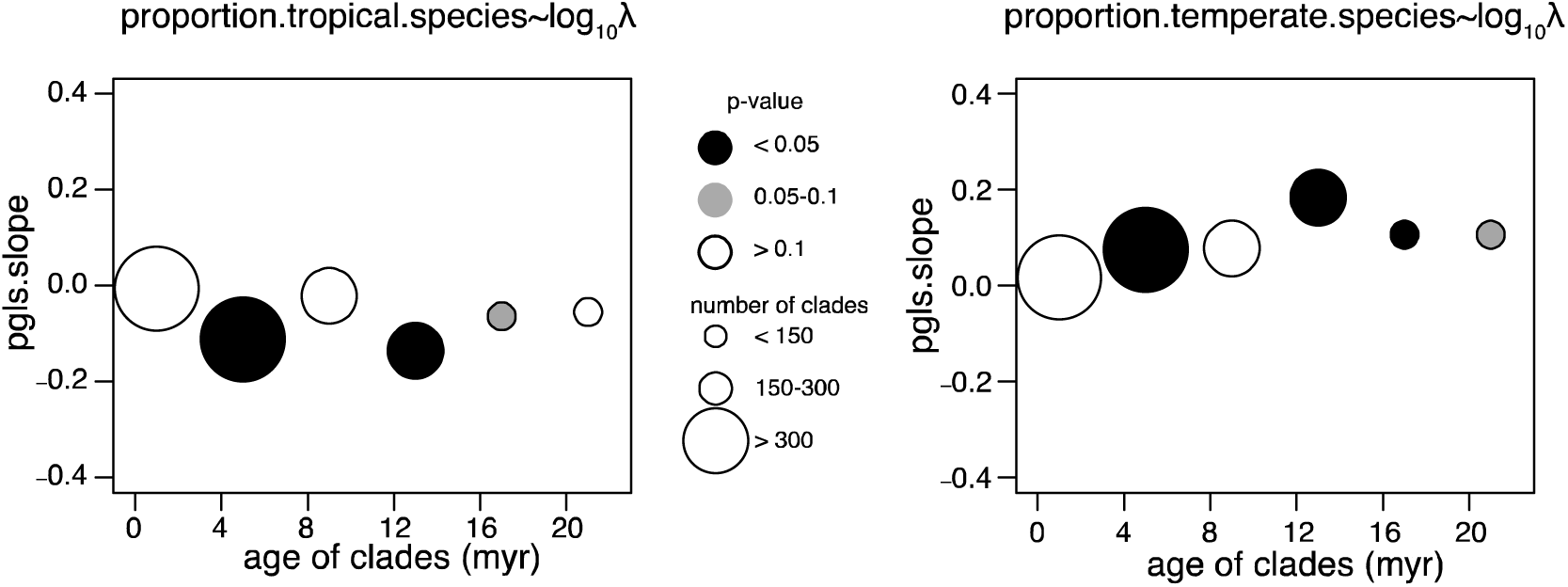
Temperate clades have higher speciation rates in the clade-based analysis. Correlation of a) the proportion of tropical and b) temperate species in each clade with the corresponding speciation rate (λ) estimated with RPANDA. The correlation coefficient is the phylogenetic generalised least squares (PGLS) slope. The size of the points is scaled to the number of clades in each time interval and their colours show the statistical significance of the slope.

**Figure S9.**
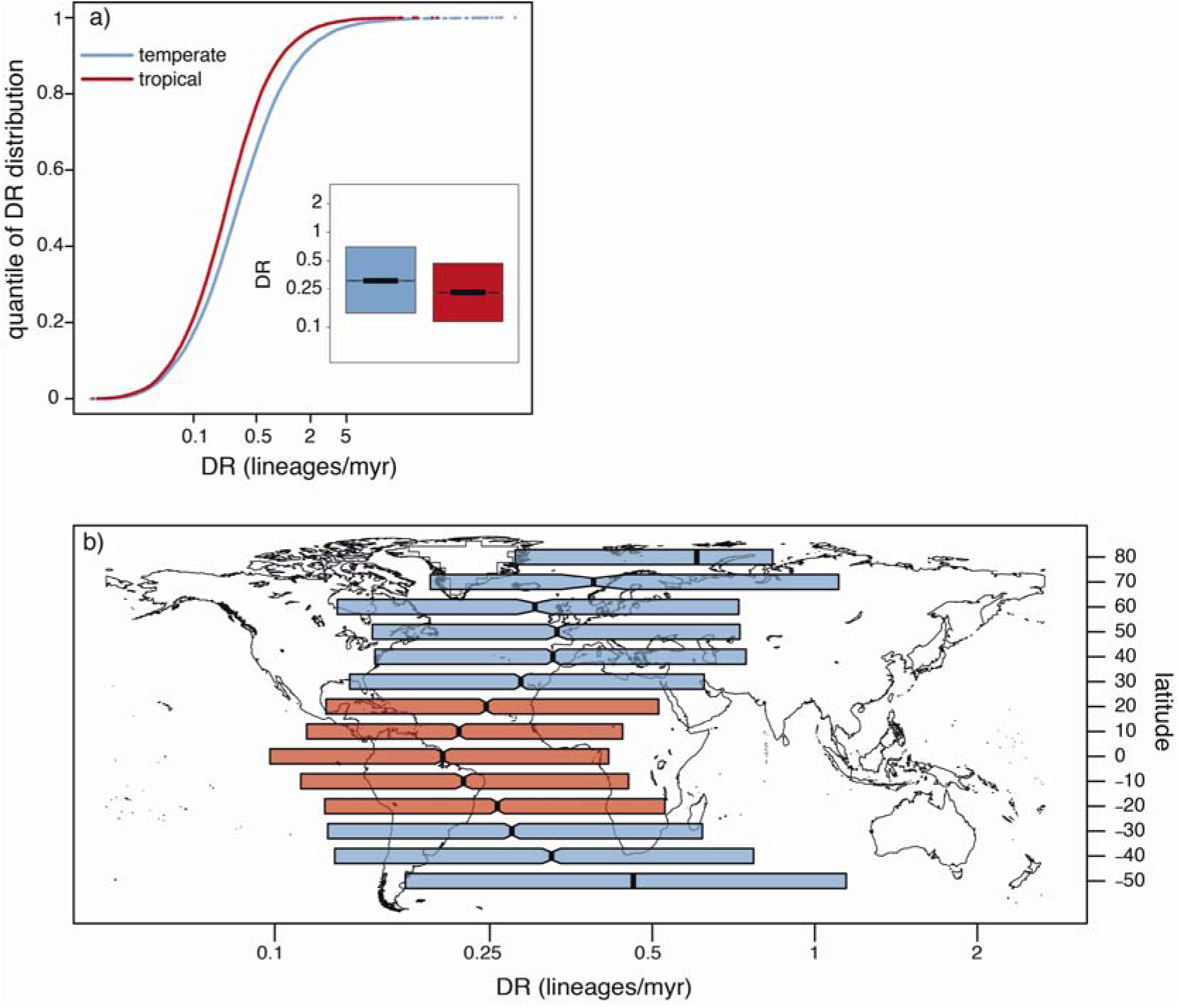
Tropical species have smaller DR values than temperate species. **a)** Rank-ordered distribution and boxplot (inset) of DR for tropical (red) and temperate (blue) species. **b)** DR grouped by latitudinal band of each species. Notches in boxplots indicate 95% confidence intervals around median, denoted by thick vertical lines and boxes span the interquartile ranges.

## Acknowledgments

We thank the Wellcome Trust (Grant Number 105602/Z/14/Z), Gatsby Charitable Foundation (Grant Number GAT2962) and Isaac Newton Trust (Grant Number 17.24r) for funding. We thank the Handling Editor, Hervé Sauquet and two anonymous reviewers for comments that improved the manuscript.

## Data accessibility

Scripts and data to reproduce the analyses will be deposited upon acceptance in Figshare

